# Central and peripheral delivery of AAV9-SMN target different pathomechanisms in a mouse model of spinal muscular atrophy

**DOI:** 10.1101/2021.11.08.467795

**Authors:** Aoife Reilly, Marc-Olivier Deguise, Ariane Beauvais, Rebecca Yaworski, Simon Thebault, Daniel R. Tessier, Vincent Tabard-Cossa, Niko Hensel, Bernard L. Schneider, Rashmi Kothary

## Abstract

Spinal muscular atrophy (SMA) is a neuromuscular disease caused by loss of the *SMN1* gene. Although lower motor neurons are a primary target, there is evidence that peripheral organ defects contribute to SMA. Current SMA gene therapy uses a single, high titre intravenous bolus of AAV9-SMN resulting in impressive, yet limited amelioration of the clinical phenotype. However, risks of this treatment include liver toxicity. Intrathecal administration is under clinical trial but was interrupted due to safety concerns in a concomitant animal study. As there is no direct comparison between the different delivery strategies while avoiding high dose toxicity, we injected SMA mice with low dose scAAV9-cba-SMN either intravenously (IV) for peripheral SMN restoration or intracerebroventricularly (ICV) for CNS-focused SMN restoration. Here, IV injections restored SMN in peripheral tissues but not CNS, while ICV injections mildly increased SMN in the periphery and the CNS. Consequently, only ICV treatment rescued motor neuron degeneration. Surprisingly, both treatments resulted in an impressive rescue of survival, weight, motor function, and peripheral phenotypes including liver and pancreas pathology. Our work highlights independent contributions of peripheral organs to SMA pathology and suggests that treatments should not be restricted to the motor neuron.

**Graphical Abstract:** 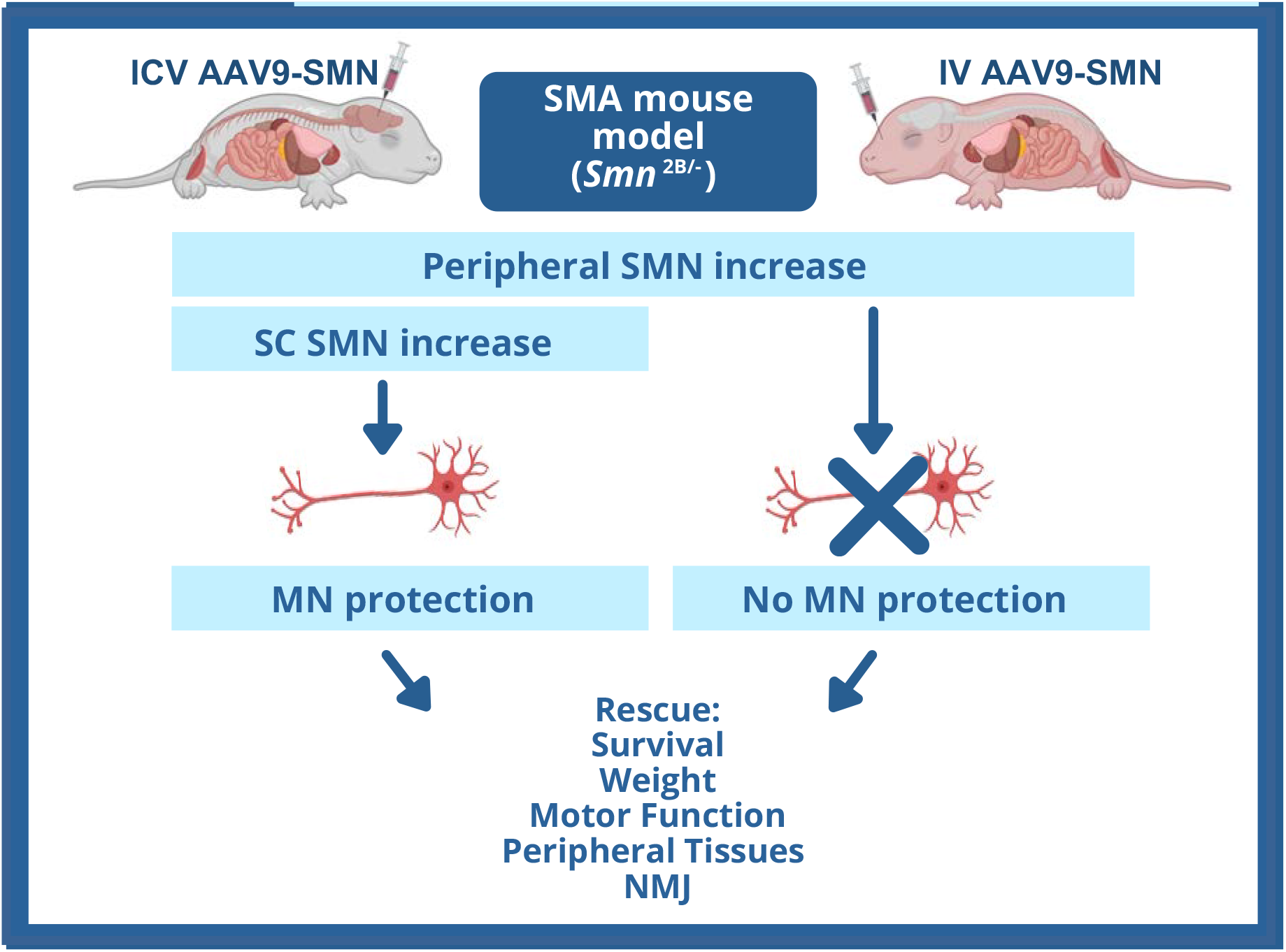

## Introduction

Spinal muscular atrophy (SMA) is a devastating childhood neurodegenerative disorder characterized by the loss of lower motor neurons and skeletal muscle atrophy. Untreated and severely affected patients suffer from proximal and progressive muscle weakness, leading to complications such as scoliosis, respiratory failure, and early death ^1^. Recent therapeutic advances, notably in an AAV9-mediated gene therapy, are bringing hope to SMA patients, but toxicity concerns of the high viral dose required demand a closer look at the therapeutic potential of low dose treatment.

SMA is caused by deletion or mutation in the *Survival of Motor Neuron 1* (*SMN1*) gene ^2^. The SMN protein product is essential and a complete SMN loss results in early embryonic lethality. SMA severity is thus mediated by a second gene, *SMN2*, which produces a limited amount of SMN protein. Recent advances in SMA therapeutics have changed the landscape of SMA disease prognosis as the use of therapeutics is becoming mainstream for SMA patients. There are currently three FDA-approved therapies available for SMA patients: nusinersen, risdiplam, and onasemnogene abeparvovec. While nusinersen and risdiplam act on the *SMN2* gene to increase SMN protein production, onasemnogene abeparvovec uses a blood-brain-barrier penetrant adeno-associated virus serotype 9 (AAV9) vector carrying SMN cDNA to encode for the missing SMN protein. This gene therapy treatment is administered as a one-time intravenous (IV) injection, aiming to induce long-term expression of SMN protein in the periphery as well as the CNS ^3,4^. However, there is no bioavailability data in humans so far. It is also unknown how long expression lasts in different tissues because episomal viral DNA is diluted during cell division, and permanent expression will likely be restricted to post-mitotic cells such as neurons ^5^. Limited data from similar AAV gene therapies targeting mitotic cells has confirmed transgene expression and clinical benefit for up to 6 years after vector administration ^6–8^. So far, systemic delivery has been approved for therapy. However, CNS specific routes of administration are being explored for this therapy, as are various viral concentrations ^9^.

Though targeting of motor neurons is often the focus for SMA therapy, a wide range of non-neuronal tissues are affected in SMA patients and animal models. Muscle intrinsic defects ^10–12^, cardiac defects ^13,14^, fatty acid metabolism defects ^15,16^, glucose metabolism defects ^17,18^, immune organ defects ^19–21^ and gastrointestinal dysfunction ^22,23^ have been observed in SMA patients and characterized in mouse models of the disease. In fact, the most effective experimental gene therapy treatments in SMA pre-clinical models were delivered systemically with the use of ubiquitous promoters ^4,24,25^. However, safety concerns have arisen over the high doses of AAV required to transduce the CNS when administered intravenously. AAV vectors are known to have a higher affinity for transduction of the liver compared to other tissues ^26^, causing liver damage in some onasemnogene abeparvovec treated patients ^4^, while primates treated with a high dose of a similar construct in a pre-clinical trial demonstrated liver toxicity, liver failure, and neuronal degeneration ^27^. Alternative routes of administration of this drug are also being explored, though a trial evaluating cerebrospinal fluid delivery of onasemnogene abeparvovec was suspended due to concerns with neuronal toxicity in pre-clinical studies ^9^. Further, new data has shown that long-term overexpression of SMN has neurotoxic effects and leads to motor dysfunction over time in AAV9-SMN treated mice ^28^. With serious risks associated with high dosage via both routes of delivery, it is important to understand the benefits associated with an approach using low doses comparing CNS versus peripheral delivery. This will allow for the ideal route of delivery to be determined, as well as the development of therapies using lower doses and fewer safety risks.

We used a low dose of scAAV9-cba-SMN paired with either IV or ICV delivery to generate two different patterns of viral transduction. IV delivery induced a strong SMN-expression in the periphery while omitting the spinal cord. ICV scAAV9-cba-SMN application resulted in a modest SMN increase. Strikingly, both applications led to an impressive and similar effect on survival, weight gain, and motor function, underlining the importance of peripheral SMN expression. However, while there was a similar rescue effect on peripheral phenotypes such as liver and pancreas defects, motor neurons were rescued in ICV injected mice only. Together, these results show the efficiency of SMN gene-replacement therapy with a low viral dose. Moreover, rescue mechanism varies depending on the delivery route and is different between a central and peripheral treatment strategy. Further studies are required that explore a combination of low dose delivery routes in an scAAV9-cba-SMN treatment regimen to fully exploit the different route-dependent benefits while avoiding toxic effects.

## Results

### Intracerebroventricular scAAV9-cba-SMN administration to *Smn*^*2B/-*^ mice results in a mild increase in SMN in the CNS and the periphery while intravenous injection results in restoration of SMN in peripheral tissues only

Our objective was to compare the phenotypic rescue of SMN replacement focused on the periphery with a replacement strategy primarily targeting the CNS. Therefore, we used a low dose of a blood-brain-barrier penetrant scAAV9 viral vector either administered IV or ICV to *Smn*^*2B/-*^ mice. This virus expresses SMN under the control of a chicken beta actin promoter like the commercially available SMA treatment onasemnogene abeparvovec **(Fig. 1A)**. To evaluate the tissue tropism of each application route, SMN protein levels in treated and untreated *Smn*^*2B/+*^ and *Smn*^*2B/-*^ mice were obtained by western blot to compare between the neuromuscular system compartment (spinal cord and muscle) and the liver (the major peripheral target of AAV9 in patients). Both application routes lead to increased SMN protein, demonstrating the functionality of the virus (**Fig. 1B,F)**. However, there was a difference in the tissue specific increases between routes of injection. ICV treatment produced a substantial yet mild increase of SMN protein levels in liver and spinal cord, and a trend towards an increase in muscle **(Fig. 1B-E)**, while IV treatment completely restored SMN protein to the muscle and liver, but not the spinal cord **(Fig. 1F-I)**. Not surprisingly, these results indicate that systemic scAAV9-cba-SMN administration more efficiently restores peripheral SMN levels compared to a CNS-directed approach. Importantly, systemic scAAV9-cba-SMN application at this titre omitted spinal cord transduction **(Fig. 1I)**, which could only be achieved by a CNS-directed approach. Therefore, subsequent benefits in IV treated animals are most likely not mediated by a rescue of motor neurons.

**Figure 1.**
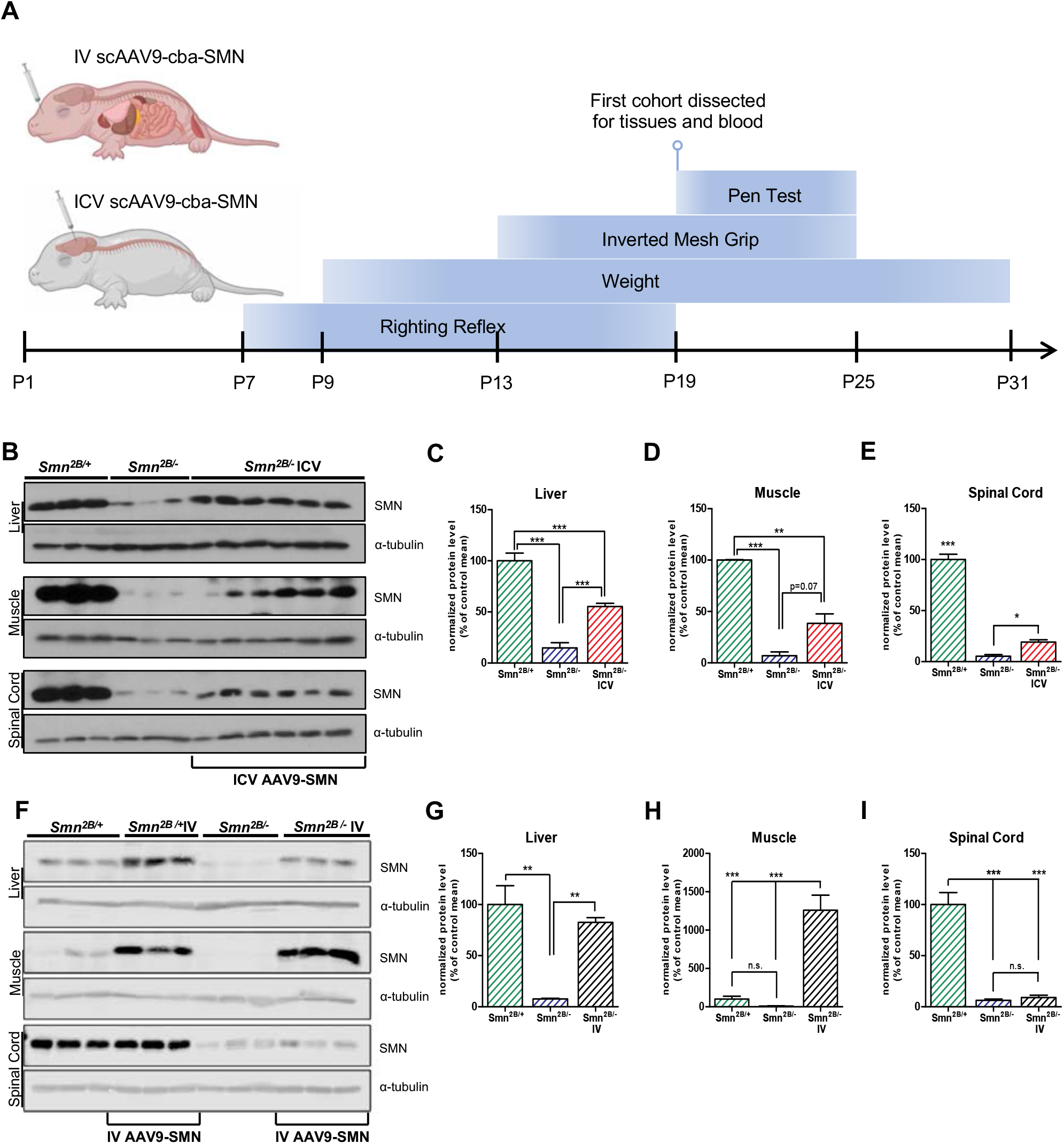
Peripheral and spinal cord SMN levels in response to ICV and IV scAAV9-cba-SMN injection. (A) Schematic representation of experimental design for AAV9-SMN treatment and evaluation of SMA-like pathophysiology. (B-E) Western blots of SMN protein levels in liver, skeletal muscle, and spinal cord following ICV injections in *Smn*^*2B/-*^ mice. (F-I) Western blots of SMN protein levels in liver, skeletal muscle, and spinal cord of IV treated *Smn*^*2B/-*^ mice. Mice were injected at P1 with 5×10^10^ vg of scAAV9-cba-SMN and tissues were dissected at P19. (n=3-6, mean ±SEM, bar graphs represent one-way ANOVA with Tukey’s post-hoc test, p ≤ 0.05 for ^*^, p ≤ 0.01 for ^**^, p ≤ 0.001 for ^***^, and n.s.= non-significant).

### Both routes of scAAV9-cba-SMN delivery significantly ameliorate SMA-like pathophysiology in *Smn*^*2B/-*^ mice with more pronounced effects after systemic administration

To compare the pathophysiological impact of peripheral SMN restoration (IV treatment) vs. partial CNS and peripheral SMN restoration (ICV treatment), we evaluated survival, body weight, and motor function. Mice were monitored every 2 days for survival and weight gain and were subjected to three motor function tests to assess muscle strength **(Fig. 2)**. Both IV and ICV treated *Smn*^*2B/-*^ mice showed a full rescue of survival within the observational period, while untreated *Smn*^*2B/-*^ mice had a median survival of 23 days **(Fig. 2A)**. Both treatments produced a partial rescue in weight gain with no differences between treatment groups **(Fig. 2B)**. IV and ICV treatment also significantly improved motor function scores. Though not significant, both treatments trended towards improved righting times. This became apparent at P19, where ICV and IV treated mice were able to immediately right themselves while *Smn*^*2B/-*^ mice generally were not **(Fig. 2C)**. In the pen test, both IV and ICV treated *Smn*^*2B/-*^ mice had significantly longer balancing times than untreated *Smn*^*2B/-*^ mice from P19 to P25 **(Fig. 2D)**. Interestingly, in the mesh grip test, times were improved in IV treated *Smn*^*2B/-*^ mice but not ICV treated mice compared to untreated *Smn*^*2B/-*^ mice **(Fig. 2E)**, suggesting greater strength recovery in distal muscles of IV treated mice. This indicates that important aspects of the SMA-like pathophysiology are influenced regardless of the locus of SMN restoration. Interestingly, IV treatment had more comprehensive impact on the motor functions emphasizing the importance of peripheral SMN restoration.

**Figure 2.**
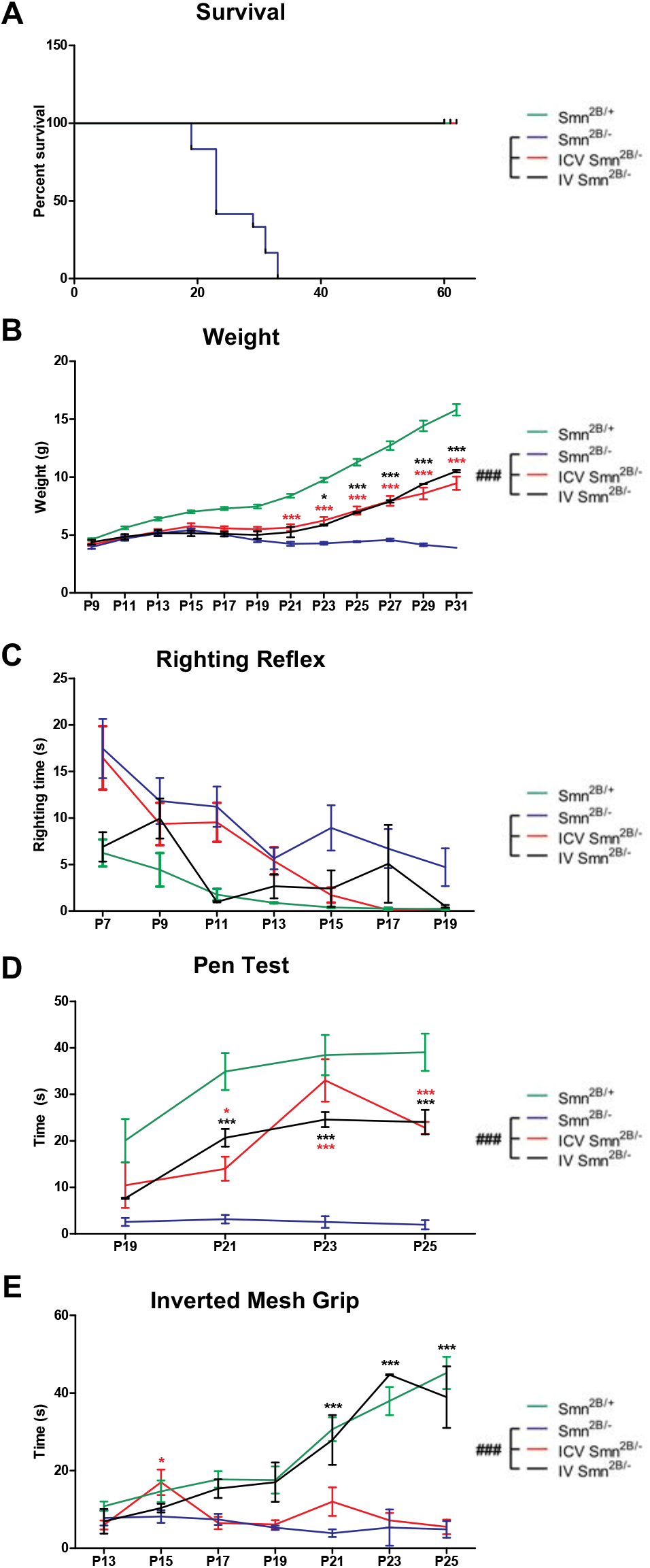
Impact of ICV and IV scAAV9-cba-SMN on SMA-like pathophysiology. (A) Kaplan Meier survival curve comparing survival of ICV and IV scAAV9-cba-SMN treated *Smn*^*2B/-*^ to saline treated *Smn*^*2B/-*^ and *Smn*^*2B/+*^ mice up to 60 days. Analysis of: (B) weight beginning from P9 to P31, (C) righting reflex from P7 to P19, (D) pen test from P19 to P25, and (E) mesh grip test from P13 to P25. Motor function and weight were evaluated every second day to compare IV treated, ICV treated, and untreated *Smn*^*2B/-*^ mice. (n=1-12, mean ±SEM, A: Kaplan-Meier survival analysis; B-E: two-way ANOVA, p≤ 0.001 for ###, with Bonferroni post-hoc test, p ≤ 0.05 for ^*^, p ≤ 0.01 for ^**^, p ≤ 0.001 for ^***^).

### CNS but not peripheral scAAV9-cba-SMN delivery partially rescues spinal cord motor neuron degeneration in *Smn*^*2B/-*^ mice

To understand the mechanisms which underly the pathophysiological changes, we next analyzed the pathological features by histology. Motor neuron number was investigated to determine the impact of IV or ICV scAAV9-cba-SMN treatment on motor neuron degeneration **(Fig. 3)**. ICV treatment partially protected against motor neuron degeneration, while there was no difference in motor neuron number between IV treated *Smn*^*2B/-*^ mice and untreated *Smn*^*2B/-*^ mice **(Fig. 3A-E)**. Additionally, we measured NfL plasma levels, which is a common outcome measure of neurodegeneration in the CNS and a candidate biomarker of neurodegeneration, disease state and therapeutic efficacy in SMA ^29^. As expected, blood plasma NfL levels were elevated in *Smn*^*2B/-*^ mice, reflecting the motor neuron degeneration **(Fig. 3F)**. Importantly, ICV treatment resulted in a reduced elevation in plasma NfL **(Fig. 3F)**. In contrast, there was no protection afforded by IV treatment as evidenced by the relatively high blood plasma NfL levels, which is in line with a lack of SMN restoration in the spinal cord. The latter result is suggestive that IV-mediated beneficial effects on pathophysiology occur despite ongoing motor neuron degeneration.

**Figure 3.**
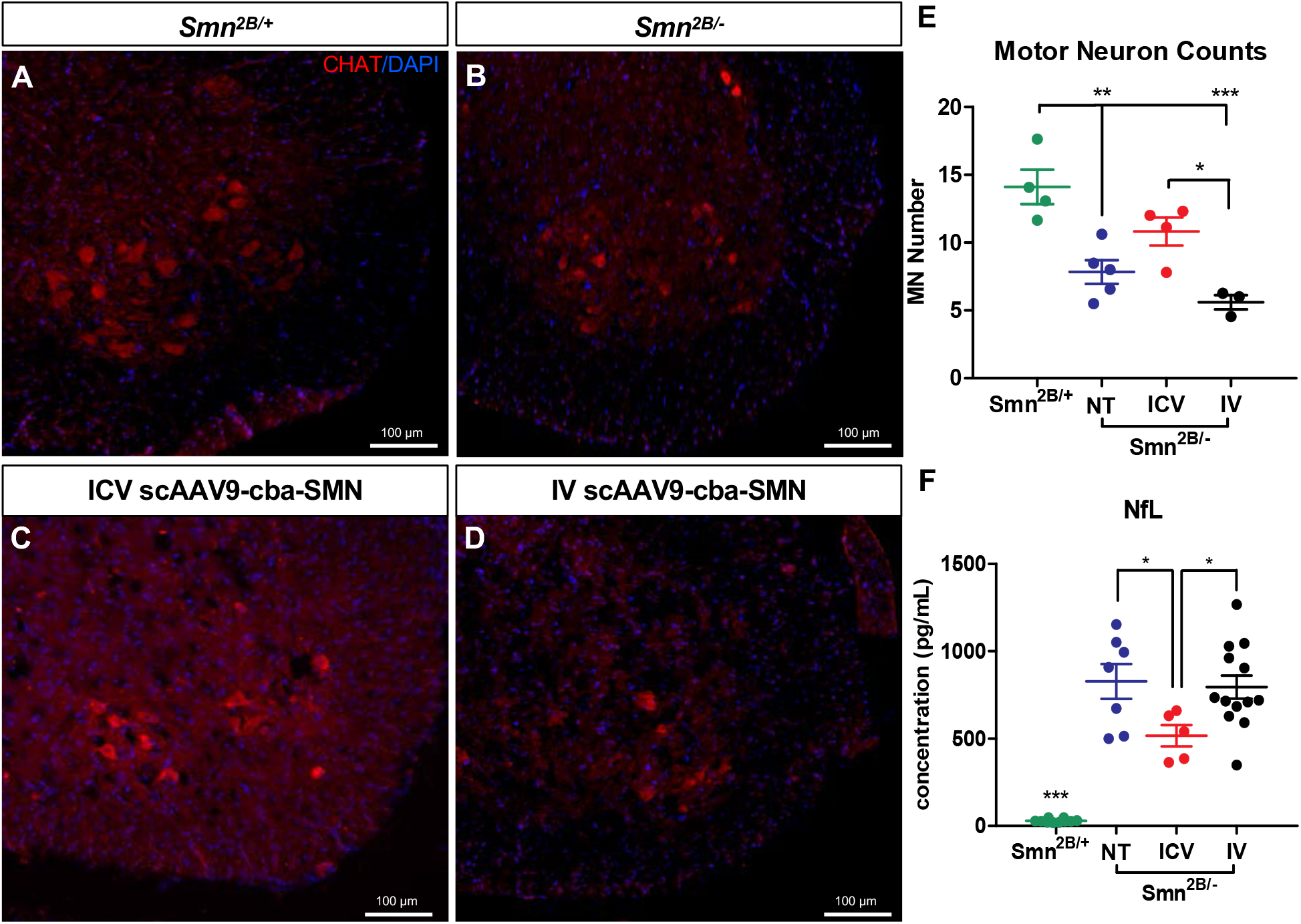
Impact of ICV and IV scAAV9-cba-SMN injection on motor neuron degeneration in *Smn*^*2B/-*^ mice. (A-D) Representative immunofluorescent images of sections of lumbar spinal cord anterior horns stained for ChAT (red) and DAPI (blue) from P19 mice. (E) Quantification of motor neuron cell body numbers. (F) Plasma neurofilament light chain (NfL) levels were assessed using single molecule array (Simoa) technology (NT=no treatment, n=4-13, mean ±SEM, one-way ANOVA with Tukey’s post-hoc test, p ≤ 0.05 for ^*^, p ≤ 0.01 for ^**^, p ≤ 0.001 for ^***^).

### CNS-directed scAAV9-cba-SMN injection in *Smn*^*2B/-*^ mice rescues neuromuscular junction pathology better than IV treatment

Next, we analysed the impact of motor neuron degeneration and/or increased muscle intrinsic SMN levels on muscle function. While there was no change in fibre size in the TA muscle by any of the scAAV9-cba-SMN injections **(Supplementary Fig. 1)**, our in-depth analyses of the NMJ pathology revealed significant differences. **(Fig. 4)**. A variety of NMJ defects are present in *Smn*^*2B/-*^ mice including abnormal endplate morphology, neurofilament accumulation, and denervation ^30^. Neurofilament accumulation and NMJ denervation were quantified to determine the degree of NMJ pathology after both routes of scAAV9-cba-SMN injection. Both IV and ICV treatments partially rescued neurofilament accumulation in the TVA of P19 *Smn*^*2B/-*^ mice, with a higher efficacy in ICV-injected animals **(Fig. 4A-D,I)**. This difference became more apparent when evaluating endplate occupancy, where only ICV injections resulted in a partial rescue **(Fig. 4E-H, J)**, which is in line with an ICV mediated rescue of motor neurons and no rescue in IV injected animals. However, the mild effect of IV-delivered scAAV9-cba-SMN on neurofilament accumulation **(Fig. 4I)** points towards a muscle-intrinsic mechanism since there is no rescue in motor neuron numbers but an increased muscle SMN expression in this group (**Fig. 3F and Fig. 1H)**.

**Fig 4.**
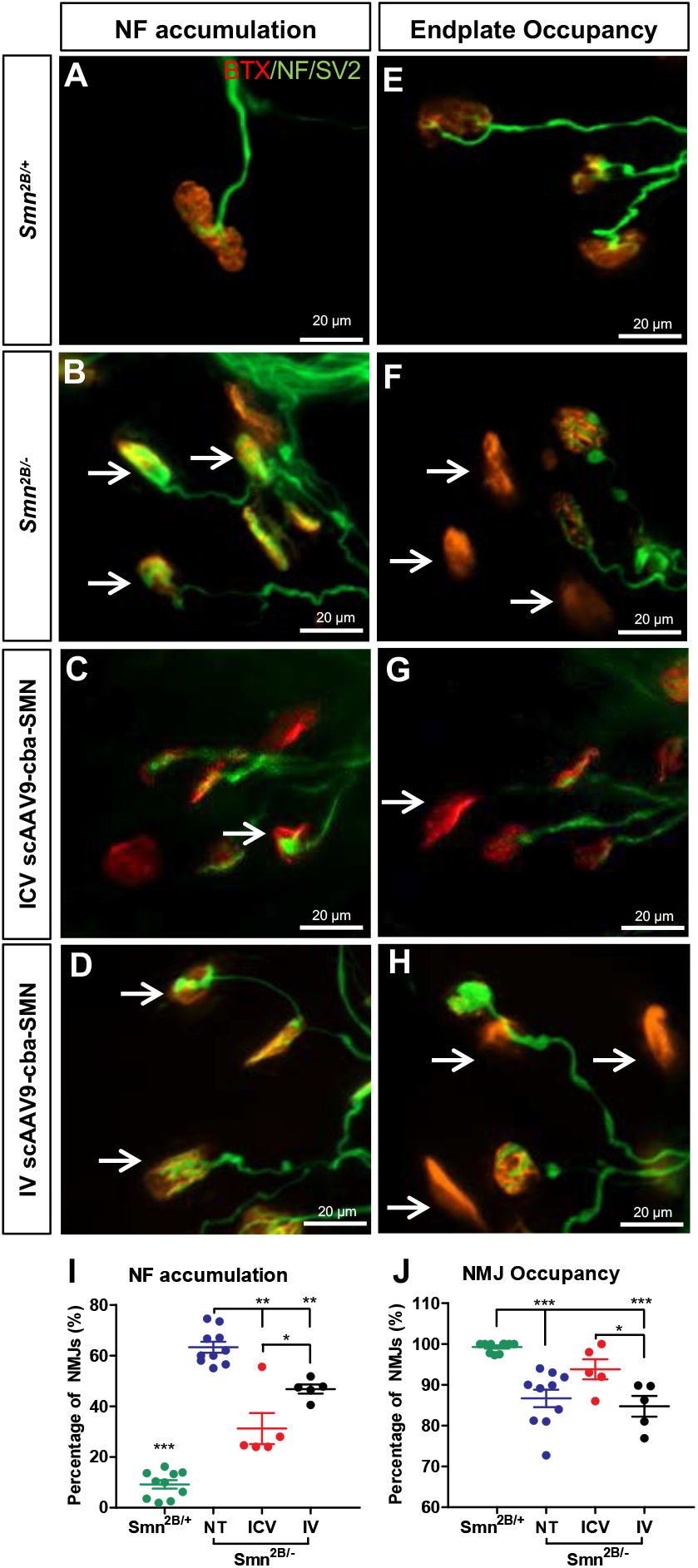
Impact of ICV and IV scAAV9-cba-SMN injection on neuromuscular junction pathology in *Smn*^*2B/-*^ mice. (A-H) Representative immunofluorescent images of *transversus abdominis* (TVA) muscle stained with bungarotoxin (red), and for neurofilament (NF) (green) and synaptic vesicle protein 2 (green) from P19 mice. Quantification of NF accumulation (I) and endplate occupancy (J) in stained NMJs to compare *Smn*^*2B/-*^ mice, *Smn*^*2B/+*^ mice, ICV treated *Smn*^*2B/-*^ mice, and IV treated *Smn*^*2B/-*^ mice. (NT=no treatment, n=5-10, B-D: arrow shows NF accumulation; F-H: arrow shows unoccupied endplate; I,J: mean ±SEM, one-way ANOVA with Tukey’s post-hoc test, p ≤ 0.05 for ^*^, p ≤ 0.01 for ^**^, p ≤ 0.001 for ^***^).

### IV and ICV scAAV9-cba-SMN treatment partially rescue peripheral organ defects in *Smn*^*2B/-*^ mice

We examined peripheral tissues to evaluate if scAAV9-cba-SMN injections affect pathological aspects that may be independent of the neuromuscular disease mechanisms. Several metabolic defects have been observed in SMA patients and mouse models of SMA. Fatty acid metabolism defects can be observed in *Smn*^*2B/-*^ mice through lipid accumulation in the liver ^15^, while glucose metabolism defects are observed through abnormal glucose homeostasis and pancreatic defects ^17^. Both IV and ICV scAAV9-cba-SMN administration prevented hepatic microvesicular steatosis in *Smn*^*2B/-*^ mice **(Fig. 5A-D)**, similar to what was previously observed with peripheral SMN restoration in this mouse model ^31^. Of note, the gastrointestinal tract of IV and ICV**C** treated mice also appeared to be healthy, unlike in untreated *Smn*^*2B/-*^ mice where it shows signs of low motility and malfunction (data not shown). Moreover, *Smn*^*2B/-*^ mice pancreata show a higher percentage of glucagon-producing alpha cells compared to insulin-producing beta cells and both treatments resulted in a partial rescue in the percentage of alpha cells within the islets compared to *Smn*^*2B/-*^ mice **(Fig. 5F-J)**. Accordingly, IV as well as ICV scAAV9-cba-SMN injections restored blood glucose levels to normal in *Smn*^*2B/-*^ mice **(Fig. 5E)**. In summary, ICV and IV injections resulted in the same degree of peripheral rescue, which is in line with the overall increase of SMN in the periphery in response to both application routes.

**Fig 5.**
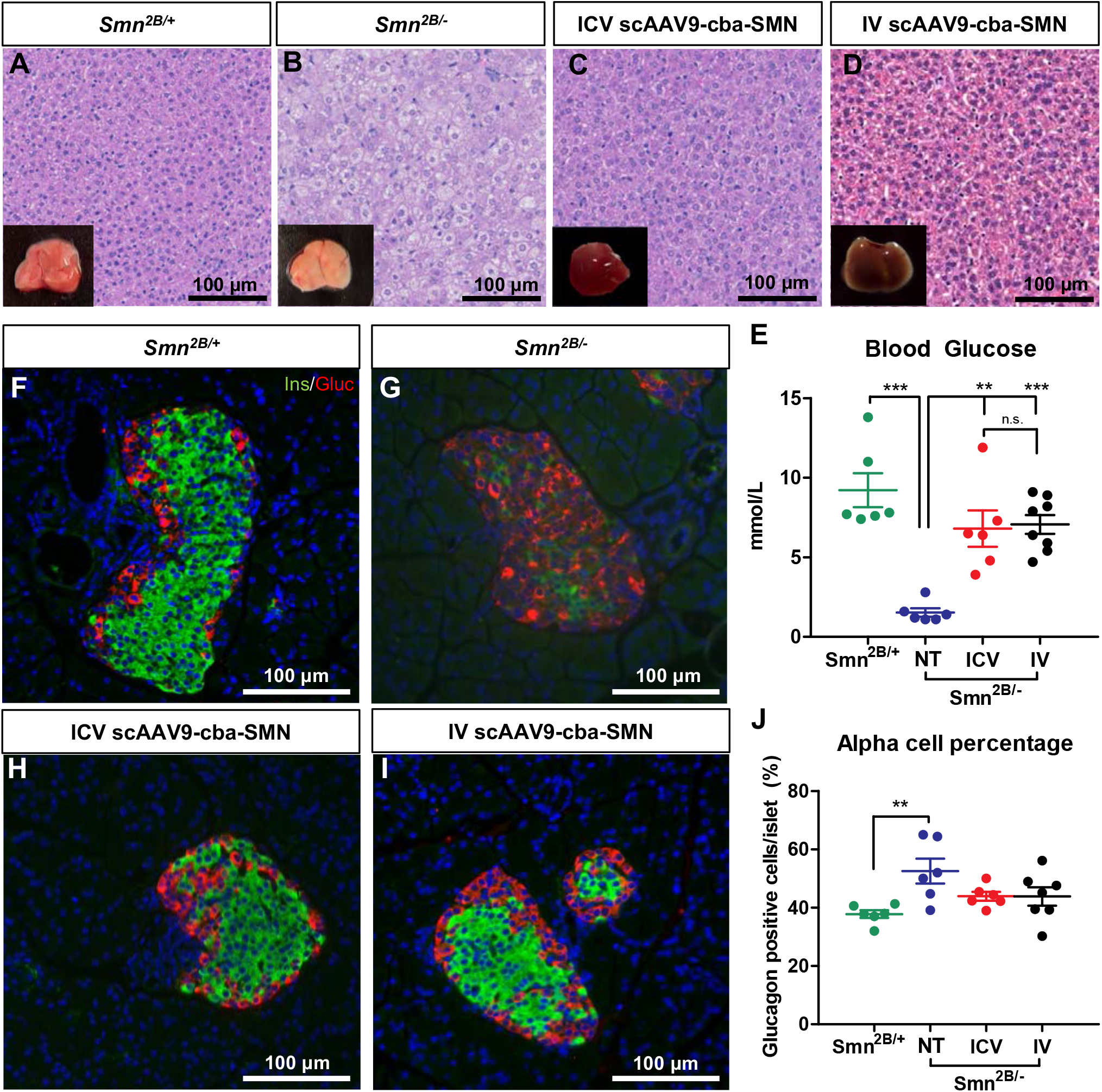
Impact of ICV and IV scAAV9-cba-SMN injection on peripheral organ defects in *Smn*^*2B/-*^ mice. (A-D) Representative images of H&E stained liver sections from P19 mice. Insets show overall appearance of the liver upon dissection. (F-I) Representative immunofluorescent images of sections of pancreatic islets stained for glucagon (red) and insulin (green) from P19 mice. (J) Fraction of glucagon-positive alpha cells related to total number of pancreatic islet cells from *Smn*^*2B/-*^ mice, *Smn*^*2B/+*^ mice, ICV treated *Smn*^*2B/-*^ mice, and IV treated *Smn*^*2B/*-^ mice. (E) Blood glucose levels from P19 mice (NT=no treatment, n=6-8, mean ±SEM, one-way ANOVA with Tukey’s post-hoc test, p ≤ 0.05 for ^*^, p ≤ 0.01 for ^**^, p ≤ 0.001 for ^***^).

## Discussion

Here, we evaluated the beneficial effects of a low dose scAAV9-cba-SMN injection in *Smn*^*2B/-*^ mice, either ICV or IV, to compare a CNS-centred SMN restoration to a peripheral-centred SMN restoration. ICV injection produced a mild increase in SMN levels within the CNS as well as in peripheral tissues of these mice, although SMN was not fully restored to wild type levels. However, the mild increase in SMN was sufficient to produce a partial rescue of neuronal degeneration and a robust rescue of the NMJ pathology, resulting in a significant amelioration of survival, weight and motor function. IV injection of the scAAV9-cba-SMN completely omitted transduction of the CNS, but fully restored SMN levels in peripheral tissues. This was surprising because of the blood-brain-barrier penetrant properties of the AAV serotype. However, this enabled us to evaluate the contribution of a peripheral SMN reduction to the overall SMA-like pathology. The significant rescue observed in weight, survival, motor function, and metabolic defects in IV-treated animals was therefore likely a result of increased SMN in peripheral tissues. In accordance with a lack of central SMN restoration, there was no rescue of motor neuron degeneration. The modest rescue of NMJ neurofilament accumulation in IV-injected mice was possibly due to muscle-intrinsic SMN restoration. Both application routes showed the same degree of rescue of liver and pancreatic phenotypes. Overall, these results, summarized in **Figure 6**, emphasize the importance of SMN restoration to the peripheral organs and suggests further investigation into the motor neuron independent mechanisms contributing to weight, survival, motor function, and metabolic defects in SMA.

**Fig 6.**
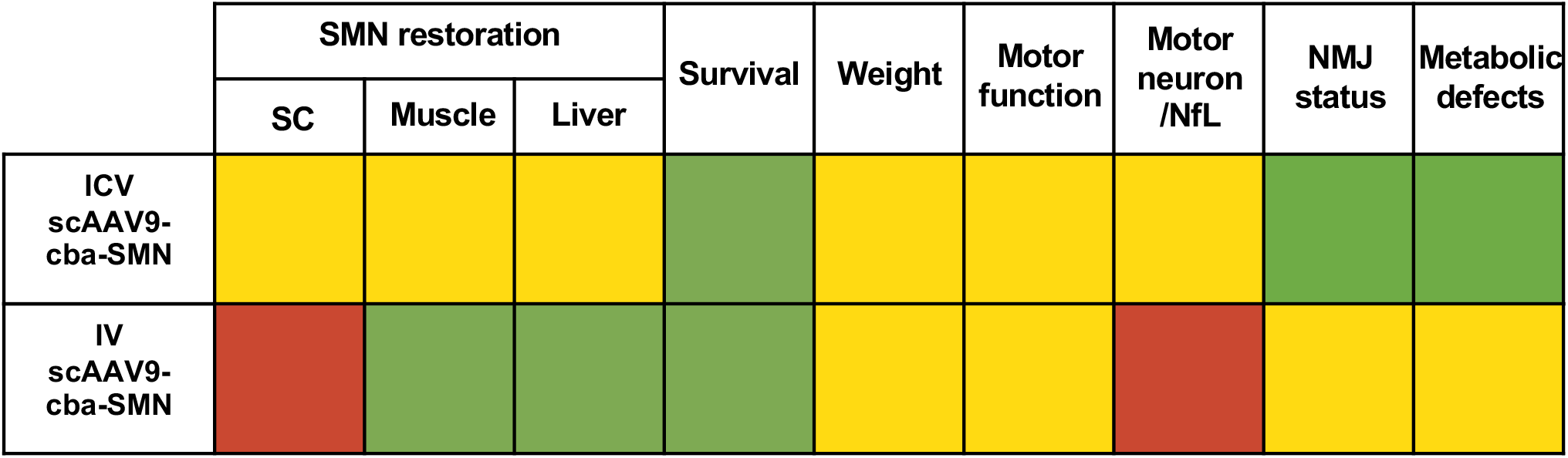
Graphical summary of the rescue of SMA-like pathology after ICV and IV scAAV9-cba-SMN injection of *Smn*^*2B/-*^ mice. Results are classified according to a colour code, where red indicates no rescue, yellow indicates some rescue, and green indicates a full rescue.

The impact of the loss of SMN protein on non-neuronal tissues is yet to be described in detail, but a variety of tissues are known to be affected in SMA patients as well as pre-clinical models of the disease ^32^. With the increasing use of SMN replacement therapy, it is important to understand the independent contributions of peripheral tissues to disease as well as the SMN protein requirements for these tissues. Our results agree with other published data that demonstrate the importance of treating the peripheral organs to achieve a full rescue. Knockdown of SMN protein in motor neurons was shown to cause an SMA-like phenotype in mice that was milder than that produced by ubiquitous knockdown ^33^, while pre-clinical nusinersen trials demonstrated 25-fold greater survival in mice receiving systemic treatment compared to CNS-restricted treatment ^34^.

A recent pre-clinical study demonstrated that AAV9-SMN gene therapy in SMNΔ7 mice restricted to neurons does not rescue the SMA phenotype in SMA mice, while ubiquitous expression does ^35^. Interestingly, this latter report demonstrated a mild increase in SMN within the spinal cord after IV injection of 4.5 × 10^10^ vg/mouse, a dose similar to our study. However, it is important to note that a different promoter, the phosphoglycerate kinase gene promoter, was used and that experiments on SMA mice using the cba promoter typically use a dose of at least 1×10^11^ vg/mouse to achieve transduction of the CNS ^25,36^. Our results emphasize the importance of targeting the peripheral organs when treating SMA and demonstrate the independent contributions of these tissues to SMA disease. This is also emphasized by clinical data, where patients treated with the CNS-specific treatment nusinersen can often remain severely disabled despite improvements in motor function, requiring ventilatory or nutritional support ^37,38^.

We were surprised to observe a slight rescue of NMJ neurofilament accumulation in our IV-injected mice, despite no restoration of SMN protein to the spinal cord in this group. This finding hinted towards possible muscle-intrinsic mechanisms of NMJ pathology, as IV injection fully restored SMN protein to muscle. However, studies have demonstrated a rescue of survival in SMA mice after muscle-specific expression of SMN, but no effect on NMJ pathology or synaptic function ^39,40^. NMJ pathology is instead likely related to SMN protein levels in motor neurons, and a consequence of motor neuron dysfunction ^41^. Interestingly however, motor neuron specific restoration of SMN in SMNΔ7 mice partially rescued endplate size and denervation, but not neurofilament accumulation ^42^. This agrees with our results that neurofilament accumulation may not be fully determined by the motor neuron, but also by other cells. Though neurofilament accumulation in neurodegenerative diseases is thought to be caused by hyper-phosphorylation of the protein within neurons ^43^, it is possible that terminal Schwann cells may have an effect on neurofilament phosphorylation and axonal transport ^44^. One explanation for the rescue of neurofilament accumulation in IV-injected mice may be that the scAAV9-cba-SMN was capable of transducing terminal Schwann cells to impact the degree of neurofilament accumulation in NMJs.

We observed a rescue in liver steatosis, with no apparent difference between the degree of rescue across the two methods of delivery. In *Smn*^*2B/-*^ mice, liver steatosis and fatty acid metabolism defects lead to several functional defects including reduced protein production, impaired hemostasis, and reduced insulin like growth factor 1 (IGF1) levels ^31^. Rescue of liver function therefore likely allowed for overall better health and survival, and a possible restoration of IGF1 levels may also have contributed to improved growth and weight gain, although this needs to be experimentally verified. We also observed a rescue in blood glucose levels and a partial rescue in pancreatic defects. *Smn*^*2B/-*^ mice demonstrate an imbalance between alpha and beta cells in the pancreatic islets, as well as glucose intolerance, hyperglucagonemia, and elevated glucose at P19 ^15,17^. Rescue of these defects likely improved survival, as these metabolic changes are associated with higher mortality and sudden cardiac death ^45^. Further, the metabolic rescue observed in IV-treated *Smn*^*2B/-*^ mice provides important evidence that the metabolic defects in SMA are motor neuron independent.

Unfortunately, systemic (IV) gene therapy presents significant risks for SMA patients due to liver toxicity and is limited in real world application due to the required high viral doses in older patients. Intrathecal application has been explored to allow for a lower dose to be used, reducing the risk of liver toxicity as well as the overall cost of the drug, but significant toxicity concerns have also been flagged for this method of delivery ^28(p9)^. This treatment method also may prevent transduction of the periphery, limiting the effectiveness of the therapy. Intravenous treatment is therefore likely worth the risk, as it is becoming more and more apparent that targeting the peripheral organs is essential for complete treatment of SMA. An alternative approach could suggest pairing a low dose systemic administration with a low dose intrathecal administration, in order to target the periphery while allowing for a less potent dose to be delivered to motor neurons and avoiding the potential toxicity associated with overexpression of SMN.

As treatments extend the lives of SMA patients and continue to change the landscape and natural history of the disease, it will be important to focus on the quality of life of SMA patients. Current treatments are by no means a cure for SMA. Treated patients may begin to develop new symptoms and face obstacles that have not been faced before by untreated patients. The data herein highlight the importance of adopting a whole-body approach to SMA treatment, focusing on each specific tissue and its independent SMN requirements. As treated patients continue to age, it is likely that peripheral symptoms will become more apparent and have a larger impact on a patient’s health. Our results display that focusing on treating the peripheral organs will not only improve patients’ quality of life but also likely impact survival and overall health.

## Materials and Methods

### Animals

*Smn*^*2B/-*^ (C57BL/6J background) mice were developed by our laboratory ^46,47^ and housed at the University of Ottawa Animal Care Facility. *Smn*^*2B/-*^ mice were produced by breeding *Smn*^*+/-*^ mice (C57BL/6J) to *Smn*^*2B/2B*^ mice (C57BL/6J). The *Smn*^*2B/-*^ mice are a model of SMA and asymptomatic heterozygous *Smn*^*2B/+*^ mice are used as controls in these experiments. Animals were cared for according to the Canadian Council on Animal Care.

### scAAV9-cba-SMN treatment

The self-complementary AAV9-cba-SMN vector was produced as previously described and was titred by real time qPCR ^31^. The scAAV9-cba-SMN expresses human SMN under the control of a chicken beta actin (cba) promoter and was produced at a titre of 9.62 × 10^13^ viral genomes (vg)/mL. Mice at post-natal day 1 (P1) were administered scAAV9-cba-SMN through facial vein injection (5×10^10^ vg per pup administered in a volume of 20 μ L; four litters of 8-12 mice) or ICV injection (5×10^10^ vg per pup in a volume of 3 μ L; three litters of 6-12 mice). Two litters from each treatment were monitored for motor function, weight, and survival and sacrificed at P60. Mice were tattooed by Animal Care Facility staff around P4-6 to allow for a specific animal’s growth and motor function to be tracked over time. Mice were weighed every 2 days. Other groups were sacrificed at P19 for collection of blood and various tissues.

### Motor function tests

Three motor function tests were performed. Righting reflex test (Treat-NMD SOP D_M.2.2.002, treat-nmd.org) evaluated overall body strength, pen test (Treat-NMD SOP SMA_M.2.1.001) assessed motor balance and coordination, and mesh grip test (Treat-NMD SOP SMA_M.2.1.002) evaluated limb strength. Motor function tests were performed as reported previously ^48^. Briefly, righting reflex test was performed from P7 to P19. In this test, the mouse is placed on its back on a flat surface and the time to right itself is measured (up to a maximum of 30 sec). Pen test involves placing the mouse on a pen and recording the length of time they are able to balance. This test was performed from P19 to P25. Mesh grip test was performed from P13 to P25. This test measures the strength of a mouse’s limbs by timing their latency to fall when suspended from a mesh. Tests were performed every 2 days. Of note, tests were performed by two different evaluators using the lab’s controlled and established protocol.

### Blood collection and plasma analysis

Mice were euthanized at P19 by decapitation after anaesthesia in a CO_2_ chamber. Upon decapitation, blood was collected using Microcuvette CB 300 K2E tubes (Sarstedt, Newton, NC) coated with K2 EDTA. Samples were spun at 2 000 g using 5424R centrifuge (Eppendorf, Hamburg, Germany) for 5 min at room temperature. Forty-five μL of the plasma supernatant was then collected in a 1.5 mL Eppendorf tube and stored at -80° C. Samples were thawed on ice and stored at 4° C the day before the assay was to be performed. Samples were analyzed using the Simoa NF-Light ® assay (Quanterix, Billerica, MA) on a Simoa HD-1 analyzer to determine the concentration of neurofilament light chain (NfL) protein. Blood glucose was measured upon blood collection using a Freestyle Precision Neo meter with Freestyle Precision Blood Glucose Test Strips (Abbott, Chicago, IL). About 1 μ L of blood was extracted from the collection tube and applied to the test strip to measure blood glucose concentration.

### Tissue processing and staining

After euthanasia at P19, liver and *tibialis anterior* (TA) muscles were fixed in 1:10 dilution buffered formalin (Thermo Fisher Scientific, Waltham, MA) for 48 h at 4°C and then transferred to 70% ethanol at 4°C until processing. Pancreata were fixed in 4% paraformaldehyde (PFA) for 48 h at 4°C and then transferred to 70% ethanol at 4°C until processing. Lumbar spinal cord was fixed in 4% PFA overnight at 4°C then prepared for cryosectioning as previously described ^48^. The abdominal musculature was dissected and fixed in 4% PFA and the *transversus abdominis* (TVA) was separated from the abdominal musculature as per ^49^.

TA, liver, and pancreas samples were processed at the University of Ottawa Department of Pathology and Laboratory Medicine and embedded in wax using a LOGOS microwave hybrid tissue processor (Milestone Medical, Kalamazoo, MI). Paraffin block tissues were cut with a microtome at 3-4 µm thickness. Sections of TA and liver were stained with hematoxylin and eosin (H&E) using an XL CV5030 autostainer (Leica, Wertzler, Germany). Samples stained with H&E were scanned with a MIRAX MIDI digital slide scanner (Zeiss, Oberkochen, Germany). Images were acquired using Panoramic Viewer 1.15.4 (3DHISTECH, Budapest, Hungary) at different magnifications. Muscle fibre size was quantified using ImageJ (version 1.53). Approximately 100-200 fibers were counted in different areas of the muscle section to ensure appropriate coverage. The area of each fibre was measured to calculate a mean fibre size for each animal. Sections of pancreas were deparaffinized in 3 washes of xylene substitute Histo-Clear (National Diagnostics, Atlanta, GA) for 5 min each followed by 2 washes of a 50/50 mixture of absolute ethanol and Histo-Clear for 5 min each. Slides were gradually rehydrated in 100% (v/v), 95% (v/v), 70% (v/v), 50% (v/v), and 0% (v/v) ethanol. Slides were incubated in 0.5% Triton-X-100 (Millipore Sigma, Burlington, MA) in PBS for 5 min, washed 3× with PBS, then blocked in 20% goat serum, 0.3% Triton-X-100 in TBS for 2 h. Slides were incubated with primary antibodies for insulin and glucagon **(Supplementary Table 1)** in 2% goat serum, 0.3% Triton-X-100 in TBS overnight at 4°C and then washed 3× with PBS. Slides were incubated with secondary antibodies **(Supplementary Table 1)** in 2% goat serum, 0.3% Triton-X-100 for 1 h and then washed 3× with PBS. DAPI (1:1 000) was added to the last PBS wash, followed by the slides being mounted in Dako Fluorescent Mounting Medium (Agilent, Santa Clara, CA). Images were taken with an Axio Imager M2 microscope (Zeiss), with a 20x objective, equipped with filters suitable for DAPI/ fluorescence.

Lumbar SC was prepared for choline acetyltransferase (ChAT) staining of motor neurons as previously desribed ^48^. The number of ChAT positive motor neurons per ventral horn was recorded for ten different sections per animal, each separated by at least 100 μ m to avoid counting the same motor neuron twice. TVA muscles were dissected and stained for neurofilament and synaptic vesicle protein to visualize neuromuscular junctions (NMJs) as before with slight alterations ^49^. Briefly, the abdominal musculature was dissected and fixed in 4% PFA and the TVA was separated from the abdominal musculature under a dissection microscope. The TVA was incubated with a tetramethylrhodamine (TRITC) conjugated bungarotoxin for 30 min at room temperature. The tissue was then incubated overnight at 4°C with primary antibodies for neurofilament and synaptic vesicle protein 2 **(Supplementary Table 1)**. Tissues were then incubated in secondary antibodies **(Supplementary Table 1)**. Tissues were mounted with Dako Fluorescent Mounting Medium (Agilent) and imaged using Axio Imager M2 microscope (Zeiss) with Z-stack feature at 40x magnification. At least 40 NMJs were counted for each animal. Each NMJ was quantified as either normal or displaying neurofilament accumulation, and endplates were noted as either occupied or unoccupied.

### Western blot

Tissue processing and immunoblotting was performed as previously described with slight alterations ^19^. After euthanasia at P19, thoracic spinal cord, liver, and TA muscle were dissected, and flash frozen in Microvette CB 300 Z tubes (Sarstedt) in liquid nitrogen. Protein was extracted from frozen tissue by homogenization of tissue with RIPA lysis buffer and PMSF (Cell Signalling, Danvers, MA). Protein concentrations of samples were determined using Bradford Assay (Bio-Rad, Hercules, CA). Twenty μ g of protein was loaded onto a 12% acrylamide gel and subject to sodium dodecyl sulfate polyacrylamide gel electrophoresis. Proteins were transferred to a PVDF membrane (Immobilon-P or Immobilon-FL, Millipore, Burlington, MA) and blocked for 1 h at room temperature in 5% (w/v) milk powder in TBS-T or Odyssey blocking buffer (Li-Cor, Lincoln, NE). See **Supplementary Table 1** for primary and secondary antibodies. Signals were detected with Odyssey CLx (Li-Cor) or using Pierce ECL Western Blotting Substrate (Thermo Scientific) and SuperSignal West Pico PLUS Chemiluminescent Substrate (Thermo Scientific). Densitometry of western blot bands was performed using ImageJ (v. 1.53). Raw values were normalized by geometric mean and used for subsequent housekeeping normalization (α-tubulin values) from the same blot.

### Statistical analysis

Data are presented as the mean ± standard error of the mean. One-way ANOVA with Tukey’s post-test or Two-way ANOVA with Bonferroni post-test were used to compare multiple means. Statistical tests were performed using GraphPad Prism 5. Significance was indicated by ^*^ for P≤0.05, ^**^ for P≤0.01, and ^***^ for P≤0.001. Images were blinded prior to quantification.

## Acknowledgements

We thank Dr. Ronald Booth and Adrienne Rowan from the EORLA at The Ottawa Hospital for assistance with some of the NfL assays.

## Funding

RK was supported by Muscular Dystrophy Association (USA) (grant number 575466); Muscular Dystrophy Canada; and Canadian Institutes of Health Research (CIHR) (grant number PJT-156379). AR was supported by a CNMD STAR Award from the University of Ottawa Brain and

Mind Institute. MOD was supported by Frederick Banting and Charles Best CIHR Doctoral Research Award.

## Competing Interests

RK received honoraria and travel accommodations from Roche as an invited speaker at their global and national board meetings in 2019. RK and the Ottawa Hospital Research Institute have a licensing agreement with Biogen for the *Smn*^*2B/-*^ mouse model. MOD received honoraria and travel accommodations from Biogen for speaking engagements at the SMA Summit 2018 held in Montreal, Canada and SMA Academy 2019 held in Toronto, Canada. These COI are outside the scope of this study. All other authors have no competing interests to declare.

## Author contributions

AR and RK designed research

AR, MOD, AB, RY, ST, and DRT performed experiments

BLS and VTC provided material support

AR, MOD, and NH analyzed the data

AR, NH, and RK wrote the manuscript with input from all authors

RK designed the study

---

**Supplementary Table 1.** List of antibodies used for immunofluorescence studies.

| **Method** | **Antibody** | **Species** | **Dilution** | **Company**<br>**(Catalog #)** |
| --- | --- | --- | --- | --- |
| IHC | ChAT | goat | 1:100 | Millipore (AB144P) |
| IHC | anti-goat Alexa<br>Fluor 555 | Donkey | 1:200 | Invitrogen<br>(A21432) |
| IHC | TRITC conjugated<br>bungarotoxin | Mouse | 1:1 000 | Invitrogen (T1175) |
| IHC | Neurofilament<br>(NF-M) | Mouse | 1:100 | (Developmental<br>Studies Hybridoma<br>Bank, P12839) |
| IHC | synaptic vesicle<br>glycoprotein 2A | Mouse | 1:250 | (Developmental<br>Studies Hybridoma<br>Bank, Q7L0J3) |
| IHC | Anti-mouse Alexa<br>Fluor 488 | Goat | 1:250 | Invitrogen<br>(A11001) |
| Western Blot | SMN | Mouse | 1:2 000 | BD Transduction<br>(610647) |
| Western Blot | Alpha-tubulin | Rabbit | 1:10 000 | Abcam (ab4074) |
| Western Blot | IRDye 680 anti-mouse IgG | Goat | 1:10 000 | Li-Cor (926-68070) |
| Western Blot | IRDye 800 anti-rabbit IgG | Goat | 1:10 000 | Li-Cor (926-32211) |
| Western Blot | Anti-mouse IgG<br>HRP conjugate | Goat | 1:3 000 | Bio-Rad (1706516) |
| Western Blot | Anti-rabbit IgG<br>HRP conjugate | Goat | 1:3 000 | Bio-Rad (1706515) |

**Supplementary Fig 1.**
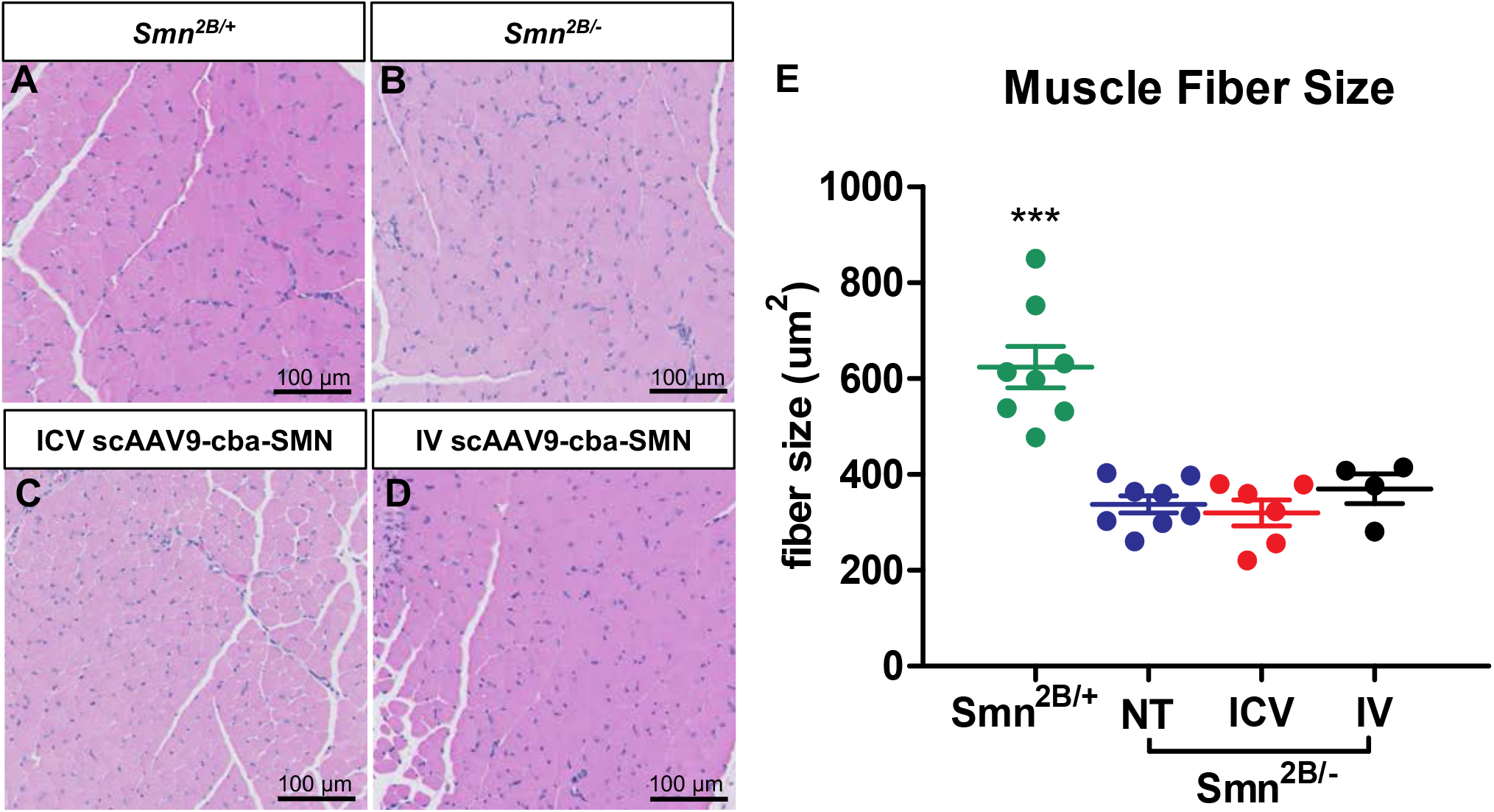
Impact of ICV and IV scAAV9-cba-SMN injection on muscle fibre size in *Smn*^*2B/-*^ mice. (A-D) Representative images of H&E-stained TA muscle sections from P19 mice. (E) Quantification of muscle fibre size area of TA muscles comparing *Smn*^*2B/-*^ mice, *Smn*^*2B/+*^ mice, ICV treated *Smn*^*2B/-*^ mice, and IV treated *Smn*^*2B/*-^ mice (NT=no treatment, n=4-8, mean SEM, one-way ANOVA with Tukey’s post-hoc test, p ≤ 0.05 for ^*^, p ≤ 0.01 for ^**^, p ≤ 0.001 for ^***^).

## References

1. Kolb SJ, Kissel JT. Spinal Muscular Atrophy. Neurol Clin. 2015;33(4):831–846. doi:10.1016/j.ncl.2015.07.004

2. Lefebvre S, Bürglen L, Reboullet S, et al. Identification and characterization of a spinal muscular atrophy-determining gene. Cell. 1995;80(1):155–165. doi:10.1016/0092-8674(95)90460-3

3. Foust KD, Nurre E, Montgomery CL, Hernandez A, Chan CM, Kaspar BK. Intravascular AAV9 preferentially targets neonatal neurons and adult astrocytes. Nat Biotechnol. 2009;27(1):59–65. doi:10.1038/nbt.1515

4. Mendell JR, Al-Zaidy S, Shell R, et al. Single-Dose Gene-Replacement Therapy for Spinal Muscular Atrophy. N Engl J Med. 2017;377(18):1713–1722. doi:10.1056/NEJMoa1706198

5. Chaytow H, Faller KME, Huang Y-T, Gillingwater TH. Spinal muscular atrophy: From approved therapies to future therapeutic targets for personalized medicine. Cell Rep Med. 2021;2(7). doi:10.1016/j.xcrm.2021.100346

6. Pennesi ME, Weleber RG, Yang P, et al. Results at 5 Years After Gene Therapy for RPE65-Deficient Retinal Dystrophy. Hum Gene Ther. 2018;29(12):1428–1437. doi:10.1089/hum.2018.014

7. Mueller C, Gernoux G, Gruntman AM, et al. 5 Year Expression and Neutrophil Defect Repair after Gene Therapy in Alpha-1 Antitrypsin Deficiency. Mol Ther J Am Soc Gene Ther. 2017;25(6):1387–1394. doi:10.1016/j.ymthe.2017.03.029

8. Rivera VM, Gao G, Grant RL, et al. Long-term pharmacologically regulated expression of erythropoietin in primates following AAV-mediated gene transfer. Blood. 2005;105(4):1424–1430. doi:10.1182/blood-2004-06-2501

9. Novartis Gene Therapies. Phase I, Open-Label, Dose Comparison Study of AVXS-101 for Sitting But Non-Ambulatory Patients With Spinal Muscular Atrophy. http://clinicaltrials.gov; 2021. Accessed July 22, 2021. https://clinicaltrials.gov/ct2/show/NCT03381729

10. Boyer JG, Murray LM, Scott K, De Repentigny Y, Renaud J-M, Kothary R. Early onset muscle weakness and disruption of muscle proteins in mouse models of spinal muscular atrophy. Skelet Muscle. 2013;3(1):24. doi:10.1186/2044-5040-3-24

11. Boyer JG, Deguise M-O, Murray LM, et al. Myogenic program dysregulation is contributory to disease pathogenesis in spinal muscular atrophy. Hum Mol Genet. 2014;23(16):4249–4259. doi:10.1093/hmg/ddu142

12. Bricceno KV, Martinez T, Leikina E, et al. Survival motor neuron protein deficiency impairs myotube formation by altering myogenic gene expression and focal adhesion dynamics. Hum Mol Genet. 2014;23(18):4745–4757. doi:10.1093/hmg/ddu189

13. Rudnik-Schöneborn S, Heller R, Berg C, et al. Congenital heart disease is a feature of severe infantile spinal muscular atrophy. J Med Genet. 2008;45(10):635–638. doi:10.1136/jmg.2008.057950

14. Shababi M, Habibi J, Yang HT, Vale SM, Sewell WA, Lorson CL. Cardiac defects contribute to the pathology of spinal muscular atrophy models. Hum Mol Genet. 2010;19(20):4059–4071. doi:10.1093/hmg/ddq329

15. Deguise M, Baranello G, Mastella C, et al. Abnormal fatty acid metabolism is a core component of spinal muscular atrophy. Ann Clin Transl Neurol. 2019;6(8):1519–1532. doi:10.1002/acn3.50855

16. Zolkipli Z, Sherlock M, Biggar WD, et al. Abnormal fatty acid metabolism in spinal muscular atrophy may predispose to perioperative risks. Eur J Paediatr Neurol EJPN Off J Eur Paediatr Neurol Soc. 2012;16(5):549–553. doi:10.1016/j.ejpn.2012.01.004

17. Bowerman M, Swoboda KJ, Michalski J-P, et al. Glucose metabolism and pancreatic defects in spinal muscular atrophy. Ann Neurol. 2012;72(2):256–268. doi:10.1002/ana.23582

18. Davis RH, Miller EA, Zhang RZ, Swoboda KJ. Responses to Fasting and Glucose Loading in a Cohort of Well Children with Spinal Muscular Atrophy Type II. J Pediatr. 2015;167(6):1362–1368.e1. doi:10.1016/j.jpeds.2015.09.023

19. Deguise M-O, De Repentigny Y, McFall E, Auclair N, Sad S, Kothary R. Immune dysregulation may contribute to disease pathogenesis in spinal muscular atrophy mice. Hum Mol Genet. 2017;26(4):801–819. doi:10.1093/hmg/ddw434

20. Khairallah M-T, Astroski J, Custer SK, Androphy EJ, Franklin CL, Lorson CL. SMN deficiency negatively impacts red pulp macrophages and spleen development in mouse models of spinal muscular atrophy. Hum Mol Genet. 2017;26(5):932–941. doi:10.1093/hmg/ddx008

21. Thomson AK, Somers E, Powis RA, et al. Survival of motor neurone protein is required for normal postnatal development of the spleen. J Anat. 2017;230(2):337–346. doi:10.1111/joa.12546

22. Gombash SE, Cowley CJ, Fitzgerald JA, et al. SMN deficiency disrupts gastrointestinal and enteric nervous system function in mice. Hum Mol Genet. 2015;24(19):5665. doi:10.1093/hmg/ddv292

23. Wang CH, Finkel RS, Bertini ES, et al. Consensus statement for standard of care in spinal muscular atrophy. J Child Neurol. 2007;22(8):1027–1049. doi:10.1177/0883073807305788

24. Dominguez E, Marais T, Chatauret N, et al. Intravenous scAAV9 delivery of a codon-optimized SMN1 sequence rescues SMA mice. Hum Mol Genet. 2011;20(4):681–693. doi:10.1093/hmg/ddq514

25. Foust KD, Wang X, McGovern VL, et al. Rescue of the spinal muscular atrophy phenotype in a mouse model by early postnatal delivery of SMN. Nat Biotechnol. 2010;28(3):271–274. doi:10.1038/nbt.1610

26. Sands MS. AAV-Mediated Liver-Directed Gene Therapy. Methods Mol Biol Clifton NJ. 2011;807:141–157. doi:10.1007/978-1-61779-370-7_6

27. Hinderer C, Katz N, Buza EL, et al. Severe Toxicity in Nonhuman Primates and Piglets Following High-Dose Intravenous Administration of an Adeno-Associated Virus Vector Expressing Human SMN. Hum Gene Ther. 2018;29(3):285–298. doi:10.1089/hum.2018.015

28. Van Alstyne M, Tattoli I, Delestrée N, et al. Gain of toxic function by long-term AAV9-mediated SMN overexpression in the sensorimotor circuit. Nat Neurosci. 2021;24(7):930–940. doi:10.1038/s41593-021-00827-3

29. Wurster CD, Steinacker P, Günther R, et al. Neurofilament light chain in serum of adolescent and adult SMA patients under treatment with nusinersen. J Neurol. 2020;267(1):36–44. doi:10.1007/s00415-019-09547-y

30. Murray LM, Comley LH, Thomson D, Parkinson N, Talbot K, Gillingwater TH. Selective vulnerability of motor neurons and dissociation of pre-and post-synaptic pathology at the neuromuscular junction in mouse models of spinal muscular atrophy. Hum Mol Genet. 2008;17(7):949–962. doi:10.1093/hmg/ddm367

31. Deguise M-O, Pileggi C, De Repentigny Y, et al. SMN Depleted Mice Offer a Robust and Rapid Onset Model of Nonalcoholic Fatty Liver Disease. Cell Mol Gastroenterol Hepatol. 2021;12(1):354–377.e3. doi:10.1016/j.jcmgh.2021.01.019

32. Deguise M-O, Kothary R. Chapter 2 - Into the unknown: Chromatin signaling in spinal muscular atrophy. In: Binda O, ed. Chromatin Signaling and Neurological Disorders. Vol 12. Translational Epigenetics. Academic Press; 2019:27–52. doi:10.1016/B978-0-12-813796-3.00002-X

33. Park G-H, Maeno-Hikichi Y, Awano T, Landmesser LT, Monani UR. Reduced Survival of Motor Neuron (SMN) Protein in Motor Neuronal Progenitors Functions Cell Autonomously to Cause Spinal Muscular Atrophy in Model Mice Expressing the Human Centromeric (SMN2) Gene. J Neurosci. 2010;30(36):12005–12019. doi:10.1523/JNEUROSCI.2208-10.2010

34. Hua Y, Sahashi K, Rigo F, et al. Peripheral SMN restoration is essential for long-term rescue of a severe spinal muscular atrophy mouse model. Nature. 2011;478(7367):123–126. doi:10.1038/nature10485

35. Besse A, Astord S, Marais T, et al. AAV9-Mediated Expression of SMN Restricted to Neurons Does Not Rescue the Spinal Muscular Atrophy Phenotype in Mice. Mol Ther. 2020;28(8):1887–1901. doi:10.1016/j.ymthe.2020.05.011

36. Meyer K, Ferraiuolo L, Schmelzer L, et al. Improving Single Injection CSF Delivery of AAV9-mediated Gene Therapy for SMA: A Dose–response Study in Mice and Nonhuman Primates. Mol Ther. 2015;23(3):477–487. doi:10.1038/mt.2014.210

37. Audic F, de la Banda MGG, Bernoux D, et al. Effects of nusinersen after one year of treatment in 123 children with SMA type 1 or 2: a French real-life observational study. Orphanet J Rare Dis. 2020;15:148. doi:10.1186/s13023-020-01414-8

38. Lavie M, Diamant N, Cahal M, et al. Nusinersen for spinal muscular atrophy type 1: Real-world respiratory experience. Pediatr Pulmonol. 2021;56(1):291–298. doi:10.1002/ppul.25140

39. Gavrilina TO, McGovern VL, Workman E, et al. Neuronal SMN expression corrects spinal muscular atrophy in severe SMA mice while muscle-specific SMN expression has no phenotypic effect. Hum Mol Genet. 2008;17(8):1063–1075. doi:10.1093/hmg/ddm379

40. Martinez TL, Kong L, Wang X, et al. Survival motor neuron protein in motor neurons determines synaptic integrity in spinal muscular atrophy. J Neurosci Off J Soc Neurosci. 2012;32(25):8703–8715. doi:10.1523/JNEUROSCI.0204-12.2012

41. Gogliotti RG, Quinlan KA, Barlow CB, Heier CR, Heckman CJ, DiDonato CJ. Motor Neuron Rescue in Spinal Muscular Atrophy Mice Demonstrates That Sensory-Motor Defects Are a Consequence, Not a Cause, of Motor Neuron Dysfunction. J Neurosci. 2012;32(11):3818–3829. doi:10.1523/JNEUROSCI.5775-11.2012

42. Paez-Colasante X, Seaberg B, Martinez TL, Kong L, Sumner CJ, Rimer M. Improvement of Neuromuscular Synaptic Phenotypes without Enhanced Survival and Motor Function in Severe Spinal Muscular Atrophy Mice Selectively Rescued in Motor Neurons. PLoS ONE. 2013;8(9):e75866. doi:10.1371/journal.pone.0075866

43. Didonna A, Opal P. The role of neurofilament aggregation in neurodegeneration: lessons from rare inherited neurological disorders. Mol Neurodegener. 2019;14(1):19. doi:10.1186/s13024-019-0318-4

44. de Waegh SM, Lee VM, Brady ST. Local modulation of neurofilament phosphorylation, axonal caliber, and slow axonal transport by myelinating Schwann cells. Cell. 1992;68(3):451–463. doi:10.1016/0092-8674(92)90183-d

45. Hess PL, Al Khalidi HR, Friedman DJ, et al. The Metabolic Syndrome and Risk of Sudden Cardiac Death: The Atherosclerosis Risk in Communities Study. J Am Heart Assoc Cardiovasc Cerebrovasc Dis. 2017;6(8):e006103. doi:10.1161/JAHA.117.006103

46. Bowerman M, Murray LM, Beauvais A, Pinheiro B, Kothary R. A critical smn threshold in mice dictates onset of an intermediate spinal muscular atrophy phenotype associated with a distinct neuromuscular junction pathology. Neuromuscul Disord NMD. 2012;22(3):263–276. doi:10.1016/j.nmd.2011.09.007

47. Eshraghi M, McFall E, Gibeault S, Kothary R. Effect of genetic background on the phenotype of the Smn2B/-mouse model of spinal muscular atrophy. Hum Mol Genet. 2016;25(20):4494–4506. doi:10.1093/hmg/ddw278

48. Deguise M-O, De Repentigny Y, Tierney A, et al. Motor transmission defects with sex differences in a new mouse model of mild spinal muscular atrophy. EBioMedicine. 2020;55:102750. doi:10.1016/j.ebiom.2020.102750

49. Murray L, Gillingwater TH, Kothary R. Dissection of the Transversus Abdominis Muscle for Whole-mount Neuromuscular Junction Analysis. JoVE J Vis Exp. 2014;(83):e51162. doi:10.3791/51162

